# Potent neutralization by a receptor binding domain monoclonal antibody with broad specificity for SARS-CoV-2 JN.1 and other variants

**DOI:** 10.1101/2024.04.27.591446

**Authors:** Michael S. Piepenbrink, Ahmed Magdy Khalil, Ana Chang, Ahmed Mostafa, Madhubanti Basu, Sanghita Sarkar, Simran Panjwani, Yaelyn H. Ha, Yao Ma, Chengjin Ye, Qian Wang, Todd J. Green, James L. Kizziah, Nathaniel B. Erdmann, Paul A. Goepfert, Lihong Liu, David D. Ho, Luis Martinez-Sobrido, Mark R. Walter, James J. Kobie

**Affiliations:** Department of Medicine, Division of Infectious Diseases, University of Alabama at Birmingham, Birmingham, Alabama, United States of America; Department of Disease Intervention and Prevention, Texas Biomedical Research Institute, San Antonio, Texas, United States of America; Department of Zoonotic Diseases, Faculty of Veterinary Medicine, Zagazig University, Zagazig, Egypt; Pacific Northwest University of Health Sciences, Yakima, Washington, United States of America; Center of Scientific Excellence for Influenza Viruses, National Research Centre, Giza, 12622, Egypt; Department of Microbiology, University of Alabama at Birmingham, Birmingham, Alabama, United States of America; Aaron Diamond AIDS Research Center, Columbia University Vagelos College of Physicians and Surgeons, New York, New York, USA

## Abstract

SARS-CoV-2 continues to be a public health burden, driven in-part by its continued antigenic diversification and resulting emergence of new variants. While increasing herd immunity, current vaccines, and therapeutics have improved outcomes for some; prophylactic and treatment interventions that are not compromised by viral evolution of the Spike protein are still needed. Using a rationally designed SARS-CoV-2 Receptor Binding Domain (RBD) – ACE2 fusion protein and differential selection process with native Omicron RBD protein, we developed a recombinant human monoclonal antibody (hmAb) from a convalescent individual following SARS-CoV-2 Omicron infection. The resulting hmAb, 1301B7 potently neutralized a wide range of SARS-CoV-2 variants including the original Wuhan and more recent Omicron JN.1 strain, as well as SARS-CoV. Structure determination of the SARS-CoV-2 EG5.1 Spike/1301B7 Fab complex by cryo-electron microscopy at 3.1Å resolution demonstrates 1301B7 contacts the ACE2 binding site of RBD exclusively through its VH1-69 heavy chain, making contacts using CDRs1-3, as well as framework region 3 (FR3). Broad specificity is achieved through 1301B7 binding to many conserved residues of Omicron variants including Y501 and H505. Consistent with its extensive binding epitope, 1301B7 is able to potently diminish viral burden in the upper and lower respiratory tract and protect mice from challenge with Omicron XBB1.5 and Omicron JN.1 viruses. These results suggest 1301B7 has broad potential to prevent or treat clinical SARS-CoV-2 infections and to guide development of RBD-based universal SARS-CoV-2 prophylactic vaccines and therapeutic approaches.

## Introduction

Severe Acute Respiratory Syndrome Corona Virus 2 (SARS-CoV-2) is the causative agent of Coronavirus Disease 2019 (COVID-19) which emerged in December 2019 in Wuhan, Hubei Province, China ^1^. Although the World Health Organization no longer considers COVID-19 a Public Health Emergency of International Concern, it continues to be a significant health threat, with over 300,000 deaths worldwide in 2023^2^. In the United States, COVID-19 has gone from the third leading cause of death in 2021 down to the tenth in 2023. The reduction in COVID-19 related deaths can likely be attributed to better treatment options and increased immunity as a result of vaccination and/or previous infection. Current COVID-19 vaccines however are sub-optimal in preventing infection, albeit a high bar for respiratory viral infections, and inducing sufficient neutralizing antibodies recognizing conserved epitopes in the viral Spike (S) glycoprotein.

With the initial emergence of the SARS-CoV-2 Omicron variant of concern (VoC) in 2021, greater vaccine breakthrough and reinfection of previously infected individuals was observed ^3^. This was in large part due to numerous mutations in the viral S protein, particularly in the receptor binding domain (RBD) which allowed for immune evasion. Each mutation has the potential of reducing the effectiveness of pre-exiting immunity afforded by previous vaccination or SARS-CoV-2 infection. JN.1, a sub-lineage of the BA.2.86 Omicron lineage, distinguished by the L455S mutation in the S protein and three non-S mutations, was the dominant circulating lineage in early 2024 ^4^. JN.1 appears to have a significant growth advantage when compared to other Omicron lineages (BA.2.86.1, XBB, and HK.3) and the L455S mutation results in considerable immune evasion ^4,5^. This emphasizes the need for updated vaccines to minimize breakthrough infections caused by JN.1 and future SARS-CoV-2 variants.

Despite reductions in SARS-CoV-2 infections; for certain individuals that are either immunocompromised or have comorbidities that increase the risk of progression of disease, the need for therapies is still essential. Although antivirals like nirmatrelvir/ritonavir and molnupiravir are available, conditions like immune system related diseases or cancer may preclude patients from receiving these drugs. Monoclonal antibodies (mAb) are generally safe with few side effects and several targeting the RBD of SARS-CoV-2 S were previously given emergency use authorization by the United States FDA. However, mAb therapies like Evusheld [tixagevimab (AZD8895) and cilgavimab (AZD1061); AstraZeneca] and Sotrovimab (S309) (Glasko Smith Kline) have reduced neutralizing activity against SARS-CoV-2 Omicron, and authorization was subsequently removed. While there are encouraging pre-clinical results for broadly neutralizing universal beta-coronavirus mAbs tolerant of the diversity of Omicron, largely targeting conserved epitopes in the S2 domain of S, the development and characterization of RBD targeting broadly neutralizing Abs that are active against emerging Omicron variants is needed as potential therapeutics and to inform future vaccine development.

Here we characterize a hmAb, 1301B7 derived from a convalescent patient following Omicron infection in the spring of 2023. This 1301B7 hmAb has sub-nanomolar binding to the RBD of the Wuhan-Hu1 strain as well as all Omicron sub- variants tested. Potent viral neutralization *in vitro* and prophylactic protection in a mouse model was also observed.

## Results

### RBD-ACE2 fusion protein-base B cell isolation

To identify and enrich for B cells that recognize epitopes within RBD that are involved with ACE2 binding, we developed a negative selection strategy using a stabilized RBD-ACE2 fusion protein complex (RBD- ACE2FP), that occludes critical ACE2 binding residues of RBD (**Figure 1A**). Using fluorescent streptavidin-tetramer probes of wild type RBDs and the RBD-ACE2FP, a differential staining strategy was utilized to selectively isolate peripheral blood memory B cells from convalescent individuals following presumed infection with SARS-CoV-2 Omicron which bind WA1 RBD and Omicron BA.2 RBD, but not the RBD-ACE2FP (**Figure 1B**). The resulting B cells were used to generate recombinant fully human IgG1 mAbs (hmAbs).

**Figure 1.**
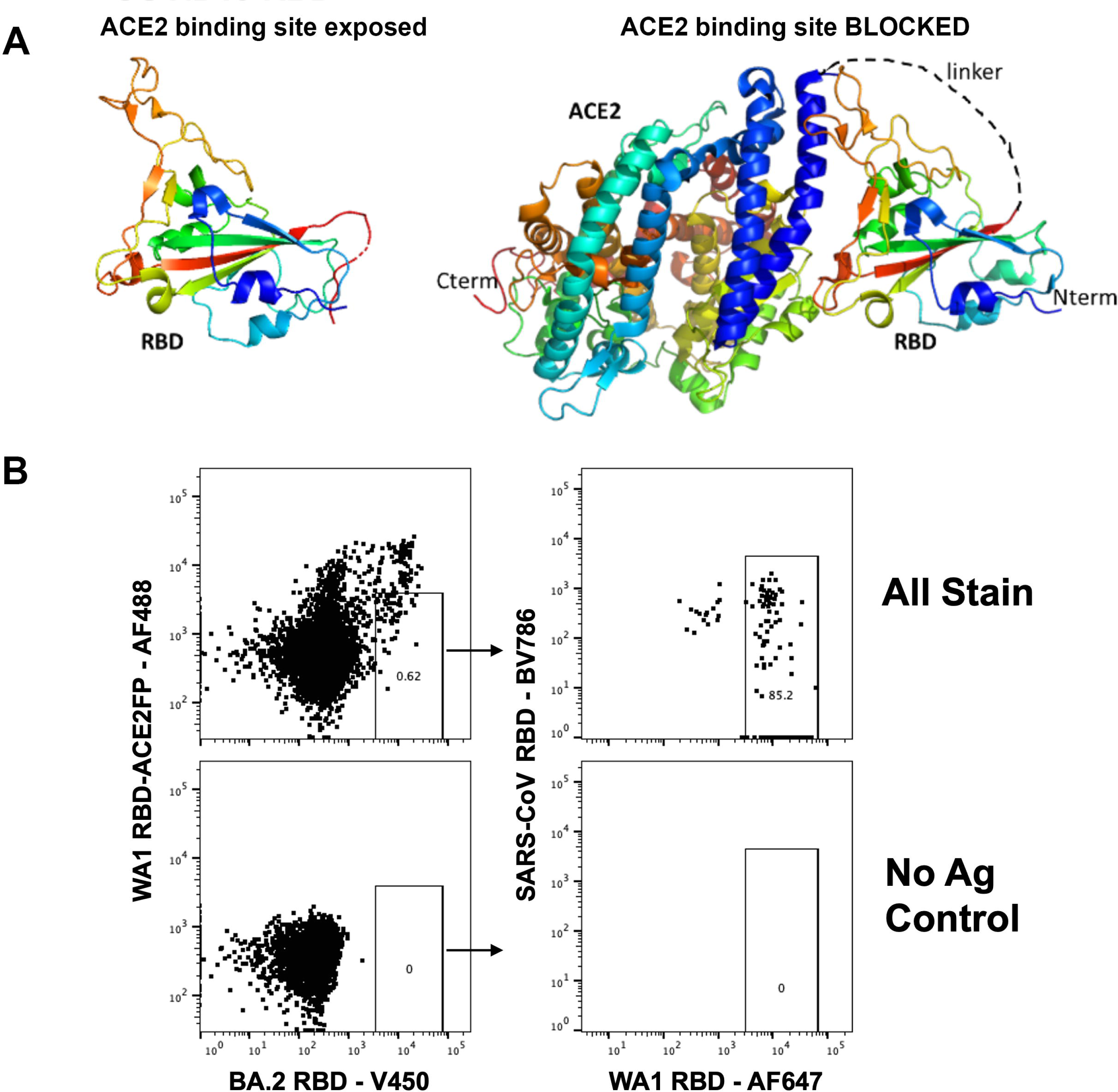
RBD-ACE2 stabilized fusion protein for isolation of B cells. (**A**) Representation of stabilized WA-1 RBD-ACE2 fusion protein. (**B**) Strategy for isolation of RBD-specific B cells by flow cytometry. Initial plot gated on live, annexin V negative, CD19+, IgD negative, HIV p24 negative, peripheral blood B cells.

### 1301B7 hmAb potently bind and neutralize SARS-CoV-2 variants

Three hmAbs with strong RBD binding emerged from initial screening. hmAbs 1300F12, 1301A8, and 1301B7 all bound RBD V483, RBD L452R T478K, and Wuhan-Hu-1 S well in the presence of 8M Urea (**Figure 2A**). 1301B7 exhibited the broadest binding profile including recognition of RBD from recent Omicron variants (XBB.1.5, XBB.1.16, EG.5.1, and FL.1) as well as binding of SARS-CoV S protein, with minimal off-target (HIV p24) binding and was advanced for further analysis. Surface plasmon resonance (SPR) indicated 1301B7 had minimal binding to RBD-ACE2FP as expected, and high affinity for WU-1 and Omicron variant RBDs (*K*= 5-100 pM), with tolerance for variability at key 416, 456, and 487 residues (**Figure 2B**). The ability of 1301B7 to compete with ACE2 for binding to SARS-CoV-2 RBD was confirmed by ELISA (**Supplemental Figure 1**). The ability of 1301B7 to bind to SARS-CoV-2 infected cells, including JN.1 was confirmed by immunofluorescence (**Supplemental Figure 2**). The functional activity of 1301B7 was tested in pseudovirus (**Figure 2C**) and live virus (**Figure 2D**) -based neutralization assays. Previously described RBD mAbs 25F9 ^6^, C68.61 ^7^, and S309 ^8^ were included as controls. 1301B7 neutralized all SARS-CoV-2 isolates tested, in addition to exhibiting neutralizing activity against SARS-CoV pseudovirus. 1301B7 had superior ability to neutralize the recent Omicron variant JN.1 compared to 25F9, C68.61, and S309. These results indicate that 1301B7 has broad and potent *in vitro binding and neutralizing* activity against SARS-CoV-2.

**Figure 2.**
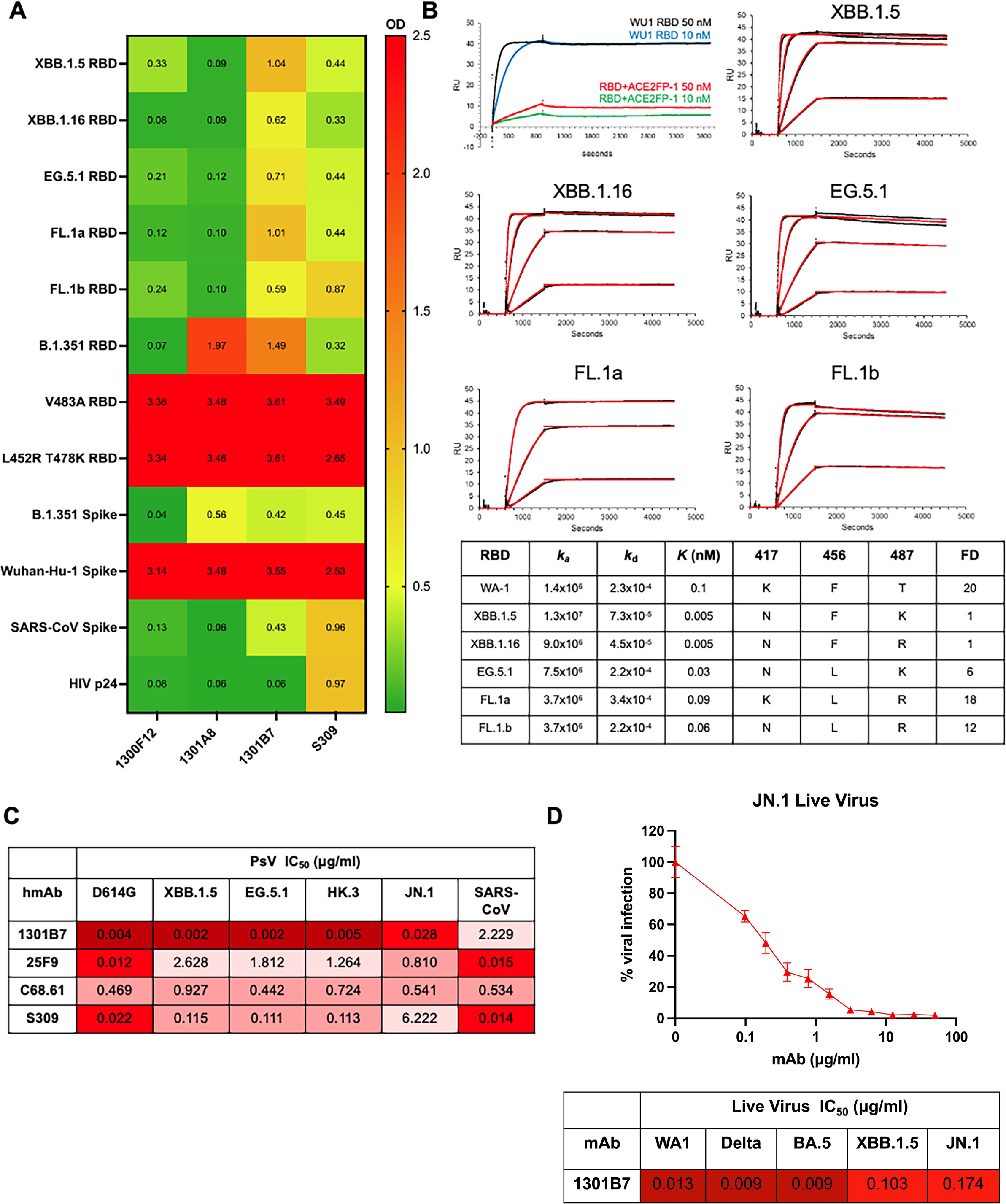
SARS-CoV-2 binding and neutralization by 1301B7 hmAb. (**A**) hmAbs were tested at 1 μg/ml in triplicate for binding to indicated protein in the presence of 8M urea by ELISA. Average optical density (OD) at 450 nM is shown. (**B**) Binding to indicated RBD was determined by SPR. Concentrations of the RBDs evaluated are 50, 12.5, 3.125, and 0.781 nM. The 50 nM concentration was not evaluated for 1301B7- FL.1 interactions. Embedded table indicates key amino acid residues and fold difference (FD) relative in *K* to XBB.1.5. hmAbs were tested at increasing concentrations in triplicate for neutralization by pseudovirus-based (**C**) or live virus (**D**) neutralization assays.

### Molecular characteristics of 1301B7 hmAb

1301B7 was isolated from an IgG1 expressing B cell and utilizes IGHV1-69 and IGLV1-40 with substantial somatic hypermutation including 20.0% amino acid mutation from germline in the heavy chain variable region, and 14.4% amino acid mutation from germline in the light chain variable region (**Table 1**). Interestingly, it has a long CDRH3 of 21 amino acids.

**Table 1:** Molecular characteristics of hmAbs.

| mAb | VH | DH | JH | Isotype | % mutation (AA) | HCDR3 length (AA) | VL | JL | % mutation (AA) | LCDR3 length (AA) |
| --- | --- | --- | --- | --- | --- | --- | --- | --- | --- | --- |
| **1301B7** | IGHV1-69 | IGHD4-17 | IGHJ6 | IgG1 | 20.0 | 21 | IGLV1-40 | IGLJ2 | 14.4 | 10 |
| **1301A8** | IGHV1-69 | IGHD1-26 | IGHJ2 | IgA1 | 16.7 | 18 | IGKV4-1 | IGKJ2 | 6.4 | 9 |
| **1300F12** | IGHV3-66 | IGHD3-16 | IGHJ6 | IgA2 | 14.4 | 16 | IGLV1-51 | IGLJ1 | 11.2 | 12 |

### Structural characterization of the 1301B7 RBD binding epitope

The cryo-EM structure of 1301B7 Fab bound to trimeric EG.5.1 Spike was determined to an overall resolution of 3.1Å (**Supplemental Figure 3**). One 1301B7 Fab binds to the single “up” RBD conformation of the Spike trimer forming a 1Fab : 1Spike trimer stoichiometry. Due to variability in orientations of the upRBD-Fab complex relative to the rest of the spike, the complex was subjected to local refinement, resulting in a 4.1Å resolution map (**Supplemental Figure 3**, **Figure 3A**). Map quality was sufficient to trace the Fab heavy chain loops and sidechains to gain insight into the 1301B7 binding epitope. The structure reveals 1301B7 predominantly (73% residue overlap with the ACE2 binding site, defined by pdbid 6m0j) contacts the ACE2 binding region of RBD (825Å^2^) exclusively through its heavy chain CDRs and FR3 region (**Figure 3B**). As previously reported for a VH1-69 anti-influenza HA receptor binding site antibody (F045-092) ^9^, 1301B7 light chain (LC) residues are ∼14Å away from the RBD. However, the space between RBD and the 1301B7 LC is filled with carbohydrate, donated from the N-linked glycosylation site (Asn100E), located within CDRH3 (**Figure 3A+3B**). The two glycans attached to CDRH3 do not form specific contacts with RBD, since the closest h-bonding atom pairs are ∼6Å away from one another.

**Figure 3.**
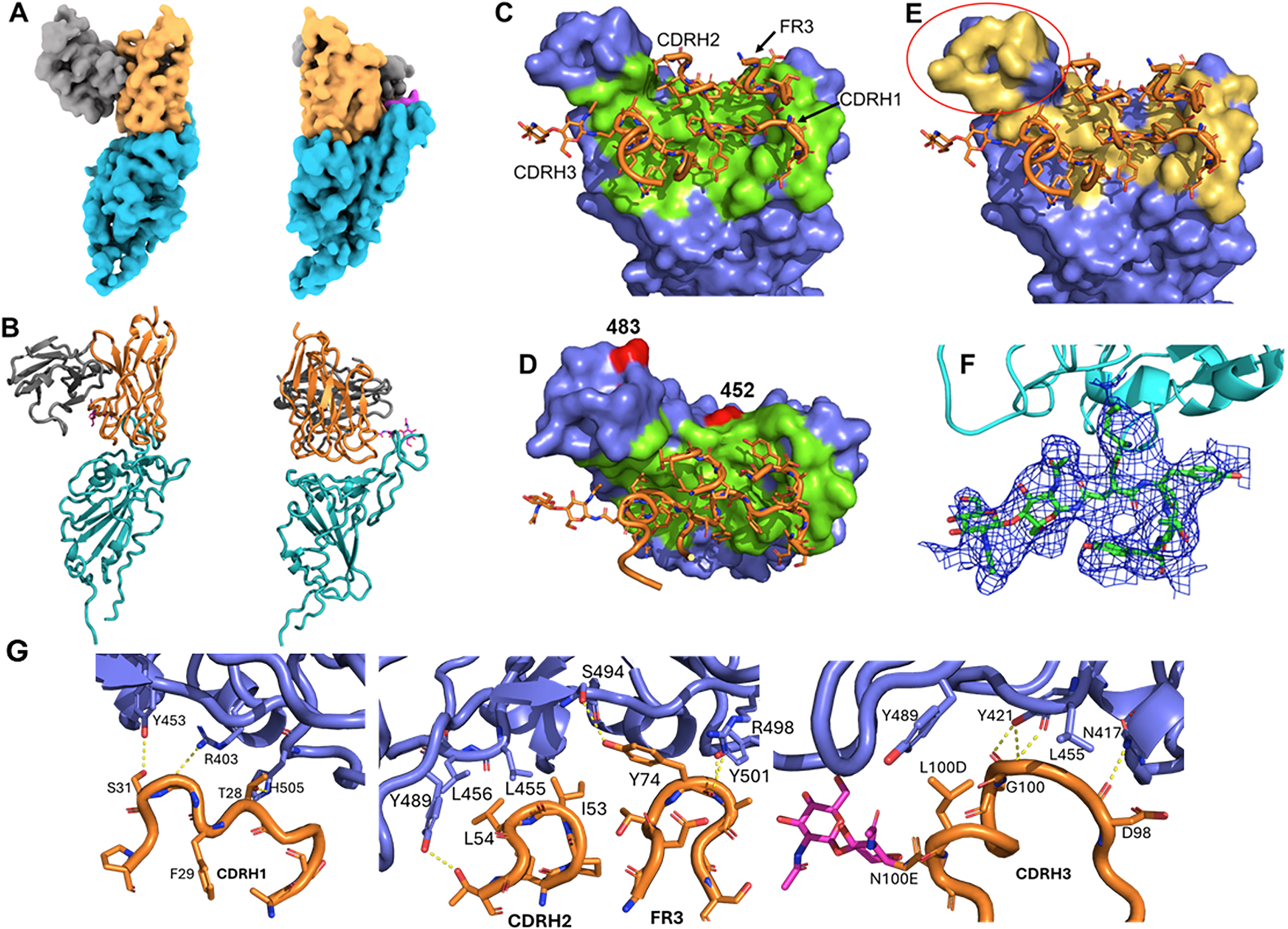
Structure of the 1301B7Fab – EG5.1 SARS-CoV-2 Spike. (**A**) Orthogonal views of the Cryo-EM density for the upRBD-1301B7 Fv complex with RBD cyan, 1301B7 heavy and light chains yellow and grey, respectively, and the glycan attached to CDRH3 is shown in magenta. (**B**) Ribbon diagram of the final model in the orientations shown in A. (**C, D**) Orthogonal views of the 1301B7 RBD binding site shown as a green surface on the EG5.1 RBD, with CDR/FR3 residues colored in green. The location of two significant JN.1 mutations L452W and A483del are shown in red. (**E**) 1301B7 CDRs/FR3 contact residues shown on the ACE2 binding surface (yellow) of RBD, showing 1301B7 does not bind to the “tip” region of RBD encircled in red on the figure 29. (**F**) Electron density from the 4.1Å local-refined map corresponding to a portion of CDRH3 and the N-linked glycan attached to Asn100E. (**G**) Molecular interactions between 1301B7 and EG5.1 RBD.

A total of 11 hydrogen bond and salt bridge interactions are observed between 1301B7 and RBD (**Figure 3D, 3E, 3F, 3G**). CDRH1 forms contacts with RBD residues R403, Y453 and H505. Notably, FR3 residue Y74 buries the greatest amount of surface area (113 Å^2^) of any residue in the interface and forms a hydrogen bond network with RBD residues S494, R498, and Y501. Signature VH1-69 hydrophobic residues ^10^ (I53, L54) at the tip of CDRH2 pack against RBD residues Y489, L456, and L455. CDRH3 buries the most surface area of any CDR into RBD and forms three h-bonds with N417, Y421 and L455. The large number of contacts in the interface are consistent with the high affinity of 1301B7 for RBD EG5.1 RBD and may partly explain its broad specificity for multiple SARS-CoV-2 variants. While the RBD N417K mutant results in a ∼20-fold drop in 1301B7 affinity, the Ab still retains sub-nanomolar binding affinity for the mutant (*K*D = 100 pM) (**Figure 2B**). Thus, 1301B7 CDRs may be able to adapt to variations in RBD sequence and structure, while maintaining high affinity and neutralizing potency.

### 1301B7 hmAb protects from lethal SARS-CoV-2 XBB1.5 infection

The K18 human ACE2 transgenic mouse model was used to determine the prophylactic activity of 1301B7 hmAb against a lethal infection with the recent SARS-CoV-2 Omicron variant XBB.1.5. Mice were treated with a single intranasal (IN) dose of 10 mg/kg or 1 mg/kg of 1301B7 6 hours prior to IN challenge with 10^5^ plaque-forming units (PFU) of XBB.1.5. All mock and isotype control hmAb treated mice exhibited substantial weight loss and had to be euthanized by day 8 post-infection (p.i.), in contrast all 1301B7 hmAb treated mice maintained their body weight and survived (**Figure 4A+B**). Treatment with 10 mg/kg of 1301B7 hmAb limited virus in the nasal turbinate, with no detectable virus at day 2 p.i. in any of the mice, and only 1 (out of 4) mouse having detectable virus in the nasal turbinate at day 4 p.i. (**Figure 4C**). No detectable virus was evident in the lungs at day 2 or 4 p.i. of mice treated with 10 mg/kg of 1301B7 hmAb (**Figure 4D**). Treatment with 1 mg/kg of 1301B7 resulted in modest, but not significant reduction in virus in nasal turbinate, however only 1 (out of 4) mouse had detectable virus in the lungs at day 2 and day 4 p.i.. These results indicate that treatment with 1301B7 hmAb protects from XBB.1.5 infection, and suggests it prevents development of viral replication in the lungs.

**Figure 4.**
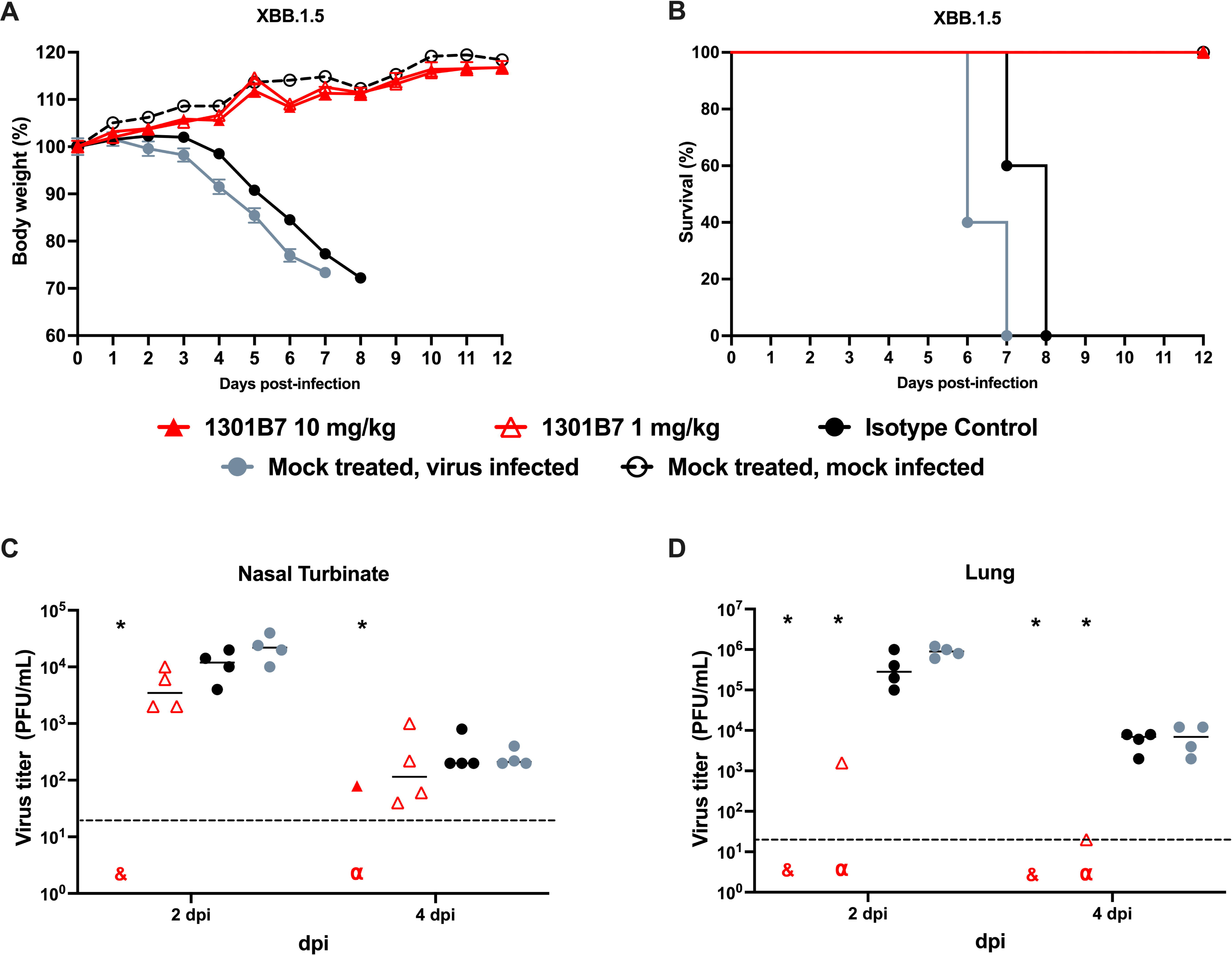
Prophylactic activity of 1301B7 hmAb against SARS-CoV-2 XBB.1.5. K18 hACE2 transgenic mice were treated i.n. with 1301B7 (1 mg/kg or 10 mg/kg) or isotype control hmAb (10 mg/kg), followed by infection with 10^5^ PFU SARS-CoV-2 XBB.1.5. Body weight (**A**) and survival (**B**) were evaluated at the indicated days post-infection (n = 5 mice/group). Mice that loss >25% of their initial body weight were humanely euthanized. Error bars represent standard error of the mean (SEM) for each group of mice. Viral titers in the nasal turbinate (**C**) and lung (**D**) at 2 and 4 days p.i. were determined by plaque assay in Vero AT cells (n = 4 mice/group/day). Symbols represent individual mice, bars indicate the mean of virus titers. Dotted lines indicate limit of detection. & indicates virus not detected in any mice from that group. α indicates virus detected in only one mouse from that group. * indicates p<0.05 as compared to isotype control hmAb as determined by t-test.

### 1301B7 hmAb protects from SARS-CoV-2 JN.1 infection

We next sought to determine the prophylactic activity of 1301B7 against JN.1 which gained predominance worldwide in late 2023. K18 hACE2 mice were IN treated with 1301B7 6 hours prior to challenge with 10^5^ PFU JN.1. Unlike XBB.1.5, untreated JN.1 infected mice had minimal weight loss, and limited mortality (**Figure 5A and B**). Mice treated with 2 mg/kg or 20 mg/kg of 1301B7 did not exhibit any weight loss or death. Viral burden was evident in the nasal turbinate and lungs of mock and isotype control hmAb treated mice (**Figure 5C and D**).

**Figure 5.**
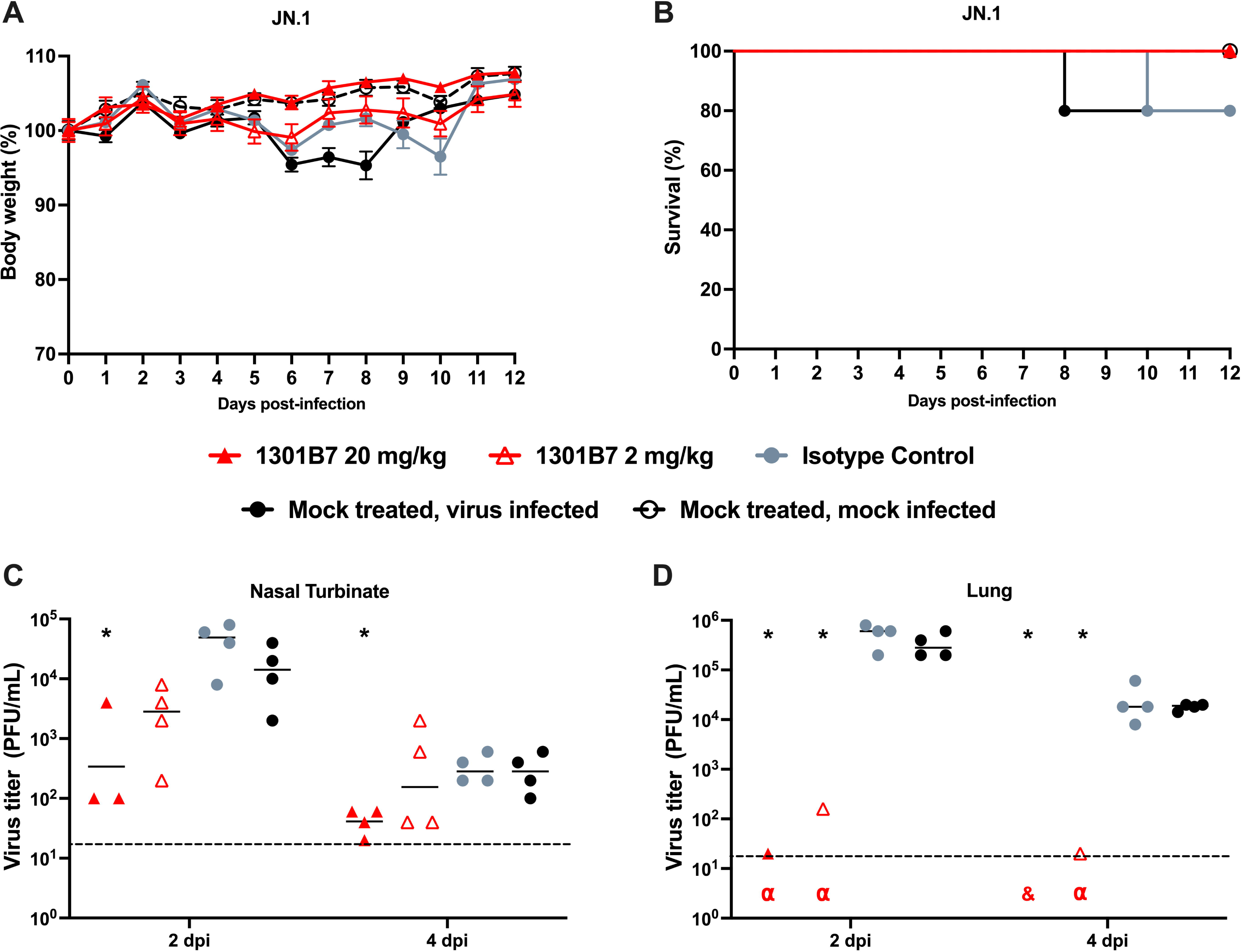
Prophylactic activity of 1301B7 hmAb against SARS-CoV-2 JN.1. K18 hACE2 transgenic mice were treated i.n. with 1301B7 (2 mg/kg or 20 mg/kg) or isotype control hmAb (20 mg/kg), followed by infection with 10^5^ PFU SARS-CoV-2 JN.1. Body weight (**A**) and survival (**B**) were evaluated at the indicated days post-infection (n = 5 mice/group). Mice that loss >25% of their initial body weight were humanely euthanized. Error bars represent standard error of the mean (SEM) for each group of mice. Viral titers in the nasal turbinate (**C**) and lung (**D**) at 2 and 4 days p.i. were determined by plaque assay in Vero AT cells (n = 4 mice/group/day). Symbols represent individual mice, bars indicate the mean of virus titers. Dotted lines indicate limit of detection. & indicates virus not detected in any mice from that group. α indicates virus detected in only one mouse for that group. * indicates p<0.05 as compared to isotype control hmAb as determined by t-test.

Mice treated with 2 mg/kg and 20 mg/kg of 1301B7 hmAb had a significant reduction (p<0.05) in nasal turbinate virus at day 2 p.i. compared to isotype control hmAb treated mice, with the significant reduction persisting at day 4 p.i. in those treated with 20 mg/kg of 1301B7. Both doses of 1301B7 resulted in significant reduction in lung virus at day 2 and day 4 p.i., with only 1 (out of 4) mouse having detectable virus in each group at day 2 p.i.. At day 4 p.i. no mice treated with 20 mg/kg of 1301B7 had detectable virus in the lungs, and only 1 (of 4) mouse treated with 2 mg/kg of 1301B7 having detectable virus in the lungs. Together these results indicate that 1301B7 has potent prophylactic activity against multiple SARS-CoV-2 Omicron variants, including the latest circulating XBB.1.5 and JN.1 strains.

## Discussion

SARS-CoV-2 is now thoroughly entrenched in humanity due to its continued ability to subvert pre-existing immunity. Thus, defining its vulnerabilities for targeted development of the highly effective next generation vaccines and therapeutics is critical to meaningful advances in reducing the impact of the virus on human health. Antibody responses to the RBD have consistently been among the most potent in neutralizing SARS-CoV-2 infection and pathogenesis, yet the activity of RBD-targeting antibodies is often negated with the development of new variants that concentrate amino acid changes in the RBD. Despite sensitivity to immune evasion, a subset of RBD mAbs are able to broadly neutralize SARS-CoV-2 variants. Here we have defined 1301B7 as a hmAb capable of broadly neutralizing all SARS-CoV-2 variants it was tested against, as well as SARS-CoV. In direct comparison, 1307B7 *in vitro* neutralizing activity was similar to other recently described RBD mAbs, and although not directly compared, its ability to protect from lethal SARS-CoV-2 Omicron XBB1.5 infection at 1 mg/kg and SARS-CoV-2 JN.1 infection at 2 mg/kg suggest it has among the most potent *in vivo* activity.

While the throughput of screening antigen-specific B cells and the B cell repertoire for neutralizing mAbs continue to increase with improving platforms, the efficiency of such campaigns will still be impacted by the initial selection process. Although we have not expressly tested the efficiency of our RBD-ACE2FP-/RBD+ differential selection approach compared to standard RBD-based positive selection, we expect it did contribute to the success in isolating 1301B7, which mimics ACE2 and potently competes with ACE2 in its binding to SARS-CoV-2 S protein. Subsequently, further application of this differential selection approach to efficiently identify and stratify RBD-specific B cells, to better define responses to various vaccine regimens, breakthrough infections, and unique patient populations is warranted and may aid in identify those B cells with broad SARS-CoV-2 protective potential.

The structure of 1301B7 bound to the EG5.1 SARS-CoV-2 Spike revealed 1301B7 binds to RBD exclusively through its heavy chain CDRs and FR3 region to generate a picomolar affinity antibody against SARS-CoV-2 variants. Consistent with its novel structure and high binding affinity, 1301B7 uses a VH1-69 heavy chain that has been shown to be elicited against the endemic viruses influenza, hepatitis, HIV-1 ^10^, as well as SARS-CoV-2 ^11^. To our knowledge, this is the first VH1-69 SARS-CoV-2 NAb that exclusively uses the heavy chain and contains an N-linked glycosylation site in CDRH3. For 1301B7, the glycan attached to CDRH3 fills the cavity between the diminutive light chain and the RBD, providing a structural scaffold to design novel broad-specificity antibodies using protein, as well as glycan-based engineering strategies. The broad specificity of 1301B7 appears to be a combination of a sub-nanomolar NAb that forms multiple h-bonds, with many other CDR residues poised to accommodate mutational variation in the RBD. Conformation of this proposed mechanism of broad specificity will be the focus of future studies.

Among SARS-CoV-2 Omicron variants there has been substantial variability in their ability to cause significant morbidity and mortality in K18 hACE2 mice ^12–14^. Observing that all untreated animals challenged with 10^5^ PFU XBB.1.5 had rapid weight loss that required euthanasia, it was unexpected that the same dose of JN.1 resulted in minimal weight loss and only 20% mortality. Nasal and lung viral titers were similar in the untreated animals between viruses, suggesting mortality was not a consequence of any gross difference in viral replication dynamics, and further determination if aspects such as inflammatory burden or secondary sites (brain, gut) may differ between the variants.

The ability of 1301B7 to limit viral titers to be below detection limit in many of the mice is encouraging and may suggest the potential to limit transmission which warrants direct assessment. It also indicates that the potential for optimal dosing of 1301B7 to mediate sterilizing protection from SARS-CoV-2 exposure should be thoroughly assessed. As we have previously demonstrated the benefits of direct respiratory delivery of mAbs for treatment of SARS-CoV-2 infection in mice, hamsters, and rhesus macaques ^15–17^, extending the utility of this approach here to recent SARS-CoV-2 variants, including JN.1, furthers supports its advancement for clinical prophylaxis and treatment of SARS-CoV-2 infections.

## Materials and Methods

### Ethics statement

All procedures and methods involving human samples were approved by the Institutional Review Board for Human Use at the University of Alabama at Birmingham (IRB-160125005). Written or oral informed consent was obtained from the participants. All experiments were performed in accordance with relevant guidelines and regulations.

### Biosafety

All *in vitro* and *in vivo* experiments with live SARS-CoV-2 were conducted in appropriate biosafety level (BSL) 3 and animal BSL3 (ABSL3) laboratories at Texas Biomedical Research Institute Biosafety (BSC), recombinant DNA (RDC), and Animal Care and Use (IACUC) committees.

### Cells and viruses

Vero E6 cells (BEI Resources NR-54970) expressing hACE2 and TMPRSS2 (Vero AT cells) were grown in Dulbecco’s modified Eagle’s medium (DMEM; Corning, Mediatech, Inc. Durham, NC, USA) supplemented with 5% fetal bovine serum (FBS, VWR) and 1% PSG (penicillin, 100 units/mL; streptomycin, 100 μg/mL; L-glutamine, 2 mM, Corning, Mediatech, Inc.) at 37 °C with 5% CO_2_. For SARS-CoV-2 infection, Vero AT cells were maintained in post-infection medium (DMEM supplemented with 2% FBS and 1% PSG) and incubated at 37 °C with 5% CO_2_. SARS-CoV-2 WA1 was obtained from BEI Resources (NR-52281). SARS-CoV-2 Omicron BA.5 (NR-58620), XBB1.5 (NR-59105), and JN.1 (NR-59694) were generously provided by Dr. Clint Florence, Program Officer, Office of Biodefense, Research Resources, and Translational Research at NIH/NIAID/DMID/OBRRTR/RRS.

### B cell isolation and hmAb generation

Peripheral blood was collected at the University of Alabama at Birmingham from adult convalescent patients approximately one month following PCR confirmed infection with SARS-CoV-2 in 2023. Peripheral blood mononuclear cells (PBMC) were isolated by density gradient centrifugation and cryopreserved. RBD-ACE2FP consists of RBD (Wuhan-1) residues 333-518 and ACE2 residues 20-598, joined by a 19 amino acid linker. RBD-ACE2FP was confirmed to have the correct size by SDS-PAGE and reactivity profile by SPR. Biotinylated RBD and RBD-ACE2FP proteins were used to create fluorescent streptavidin tetramers as previously described ^16^. Cryopreserved cells were thawed and then stained for flow cytometry similar as previously described [18], using, HIV p24-PE, RBD-ACE2FP-SA- AlexaFluor488, WA1 RBD-SA-AlexaFluor647, BA.2 RBD-SA-BV450, SARS-CoV1 RBD- SA-BV786, CD3-BV510 (OKT3, Biolegend), CD4-BV510 (HI30, Biolegend), CD14-BV510 (63D3, Biolegend), anti- CD19-APC-Cy7 (SJ25C1, BD Biosciences), IgD-PE- Cy7 (IA6-2, Biolegend), Annexin V-PerCP-Cy5.5 (Biolegend), and Live/Dead aqua (Molecular Probes). Single B cells were sorted using a FACSMelody (BD Biosciences) into 96-well PCR plates containing 4 μl of lysis buffer as previously described [37]. Plates were immediately frozen at −80°C after sorting until thawed for reverse transcription and nested PCR performed for IgH, Igλ, and Igκ variable gene transcripts as previously described ^18,19^. Immunoglobulin sequences were analyzed by IgBlast (www.ncbi.nlm.nih.gov/igblast) and IMGT/V-QUEST (http://www.imgt.org/IMGT_vquest/vquest) to determine which sequences should lead to productive immunoglobulin, to identify the germline V(D)J gene segments with the highest identity, and to scrutinize sequence properties. Paired heavy and light chain genes were cloned into IgG1 expression vectors and were transfected into HEK293T cells and culture supernatant was concentrated using 100,000 MWCO Amicon Ultra centrifugal filters (Millipore-Sigma, Cork, Ireland), and IgG captured and eluted from Magne Protein A beads (Promega, Madison, WI) as previously described ^18,19^. 25F9 and C68.61 were previously described ^6,7^ and heavy and light chain variable regions synthesized by Twist Biosciences on reported sequences and cloned into IgG1 expression vector for production in HEK293T cell.

### Binding characterization

Recombinant proteins used include SARS-CoV-2 RBD WU1, RBD XBB.1.5, RBD XBB.1.16, RBD EG.5.1, RBD FL.1a, and RBD FL.1b produced in house (Supp Figure 1), and RBD B.1.351 (BEIR, NR-55278), RBD V483A (Acros, SPD- C52H5), RBD L452R T478K (Sino, 40592-V08H90), Spike Whuan-Hu-1 (BEIR, NR- 52308), SARS-CoV1 Spike (BEIR, NR-686), and HIV p24 (Abcam, ab43037). ELISA plates (Nunc MaxiSorp; Thermo Fisher Scientific, Rochester, NY) were coated with recombinant CoV proteins at 1_μg/ml. Purified hmAbs were diluted in PBS, added to ELISA plate and incubated for 1 h followed by 8M urea treatment for 15 minutes. Binding was detected with HRP-conjugated anti-human IgG (Jackson ImmunoResearch, West Grove, PA). SPR experiments were performed on a Biacore T200 (Cytiva) at 25°C using a running buffer consisting of 10mM HEPES, 150mM NaCl, 0.0075% P20. Kinetic binding analysis for 1301B7 was performed by capturing the hmAbs to the chip surface of CM-5 chips using a human antibody capture kit (cytiva). The binding kinetics for the interaction between hmAbs and SARS-CoV-2 Spike protein (R&D Systems, 10549-cv) was determined by injecting two to four concentrations of SARS-CoV-2 RBD (50 nM highest concentration). All SPR experiments were double referenced (e.g., sensorgram data was subtracted from a control surface and from a buffer blank injection). The control surface for all experiments consisted of the capture antibody. Sensorgrams were globally fit to a 1:1 model, without a bulk index correction, using Biacore T-200 evaluation software version 1.0.

### Neutralization

hmAbs were tested for neutralization of live SARS-CoV-2 variants as previously described ^20^. A mixture of serially diluted hmAb (starting concentration of 50 μg/ml) and 200 PFU/well of SARS-CoV-2 in post-infection media were incubated at 37°C and 5% CO_2_. After 1 h incubation, Vero AT cells (96-well plate format, 4 × 10^4^ cells/well, quadruplicates) were infected with the hmAb-virus mixture. After 1 h of viral adsorption, the mixture overlay was changed with 100 μl of post-infection media containing 1% Avicel. At 18 h p.i. (14 h p.i. for WA1 strain), infected cells were fixed with 10% neutral formalin for 24 h and immunostained with the anti-NP monoclonal antibody 1C7C7 ^20^. Virus neutralization was quantified using an ELISPOT plate reader, and the percentage of infectivity calculated using sigmoidal dose response curves. The formula to calculate percent viral infection for each concentration is given as [(Average # of plaques from each treated wells–average # of plaques from “no virus” wells)/(average # of plaques from “virus only” wells—average # of plaques from “no virus” wells)] x 100. A non-linear regression curve fit analysis over the dilution curve was performed using GraphPad Prism to calculate NT_50_. Mock-infected cells and viruses in the absence of hmAb were used as internal controls.

hmAbs were also tested using a S-pseudotyped virus (PsV) asay. Pseudotyped SARS-CoV-2 was generated using a protocol established previously ^21^. Initially, HEK293T cells were transfected with SARS-CoV-2 S-encoding plasmids using 1 mg/ml of PEI and cultured for 24 h. Following this, the transfected HEK293T cells were infected with VSV-G pseudotyped ΔG-luciferase virus (Kerafast, EH1020-PM) at a multiplicity of infection (MOI) of approximately 3 to 5. After a 2 h incubation, the cells were washed three times with complete culture medium and were subsequently maintained in fresh medium for an additional 24 h. The transfection supernatant was then harvested and clarified by centrifugation at 2,000 rpm for 10 minutes. Each viral stock was subsequently incubated with 20% I1 hybridoma (ATCC, CRL-2700) supernatant for 1 h at room temperature to neutralize contaminating VSV-G particle before measuring titers and making aliquots for storage at −80 °C until use. Pseudoviruses were subjected to titration to standardize the viral input prior to each neutralization assay. hmAbs were diluted from an initial concentration of 20 µg/ml by a factor of five in 96-well plates, in triplicates. Subsequently, 50 µL of each dilution of mAb was incubated with 50 µL of diluted pseudovirus for 1 h at 37 °C, followed by the addition of 100 µL of resuspended Vero-E6 cells at a density of 3_×_10^6^ cells/ml. Wells without mAbs (serving as ’virus alone’ controls) were included in all plates. The plates were then incubated at 37°C overnight before the quantification of luciferase activity using the Luciferase Assay System (Promega) on SoftMax Pro v7.0.2 (Molecular Devices). The reduction in luciferase activity for each mAb dose, compared with the ’virus alone’ controls, was calculated. Neutralization IC_50_ values for the mAbs were obtained by fitting the data to a nonlinear five-parameter dose-response curve using GraphPad Prism v9.2

### Cyro-EM structural analysis

DNA for the SARS-CoV-2 EG5.1 S ectodomain with 6P, furin mutations, and a 6His C-terminal tag was synthesized and placed into the pTwist-CMV expression vector (Twist Bioscience). S Trimer protein was produced by transfecting the plasmid into EXPI293 cells following manufacturer’s instructions. After 6 days, the media was harvested and dialyzed into 20 mM Tris, pH 8, 0.5 M NaCl, 5mM Imidazole. The S trimer was purified by nickel affinity chromatography (Novagen), followed by gel filtration chromatography. 1301B7Fab was produced by cleavage of the antibody with papain for 7 h. Following cleavage, the FC was removed by protein-A affinity chromatography. The Fab was concentrated/exchanged into 20 mM Hepes, pH 7.5, 50 mM NaCL and separated from uncut mab species by gel filtration chromatography.

The Cryo-EM sample was prepared by heating the S trimer for 10 minutes at 37 °C, followed by the addition of a 1.3 molar ratio of 1301B7Fab. The sample (3 μl) was immediately added to C-flat R2/1 300 mesh grids placed into an Vitrobot Mark IV at 1.5 mg/ml concentration for 30 seconds and then blotted for 4-5 seconds, prior to plunge freezing. Grids were imaged on a Glacios 2 cryo-TEM, outfitted with a Falcon 4i detector using EPU software. All cryo-EM processing, from movies to final refinement, were performed using cryoSPARC v4.4.1 ^22^. Masking and density segmentation were performed using segger ^23^, implemented in ChimeraX ^24^. Model building was performed using coot ^25^ and refined using Phenix ^26^. Ribbon diagrams were made using pymol ^27^ and chimeraX.

### K18 hACE2 transgenic mice experiments

All animal protocols involving K18 hACE2 transgenic mice were approved by the Texas Biomedical Research Institute IACUC (1718MU). Five-week-old female K18 hACE2 transgenic mice were purchased from The Jackson Laboratory and maintained in the BSL3 animal facility at Texas Biomedical Research Institute under specific pathogen-free conditions. Mice were treated with a single dose of hmAb delivered i.n. 6 h prior to viral challenge. For virus infection, mice were anesthetized following gaseous sedation in an isoflurane chamber and i.n. inoculated with viral dose of 10^5^ PFU per mouse. After viral infection, mice were monitored daily for morbidity (body weight) and mortality (survival rate) for 12 days. Mice showing a loss of more than 25% of their initial body weight were defined as reaching the experimental end point and humanely euthanized. Nasal turbinate and lungs from mock or infected animals were homogenized in 1 ml of PBS for 20 s at 7,000 rpm using a Precellys tissue homogenizer (Bertin Instruments). Tissue homogenates were centrifuged at 12,000 × g (4°C) for 5 minutes, and supernatants were collected and titrated by plaque assay and immunostaining as previously described ^28^.

### Statistical analysis

Significance was determined using GraphPad Prism, v8.0. Two-tailed t tests were applied for evaluation of the results between treatments. Significance was declared at p<0.05. For statistical analysis viral titers were log transformed and undetectable virus was set to the limit of detection.

## Acknowledgments

We are grateful for the clinical research staff that enabled this project and the assistance of the University of Alabama at Birmingham (UAB) Center for AIDS Research, the UAB Flow Cytometry & Single Cell Core Facility, UAB Multidisciplinary Molecular Interaction Core Facility, and UAB Cryo-EM Facility. We thank BEI Resources for providing reagents. We also want to thank Dr. Clint Florence, Program Officer, Office of Biodefense, Research Resources, and Translational Research at NIH/NIAID/DMID/OBRRTR/RRS for providing SARS-CoV-2 Omicron BA.5, XBB1.5 and JN.1. We are most grateful for the participation of the study volunteers.

## Author contributions

J.J.K., L.M.-S., and M.R.W. conceived the study. M.S.P., A.C., M.B., S.S., and J.J.K. isolated and screened the mAbs. M.S.P., M.B, S.S., H.H., J.J.K., S.P., M.R.W., and S.P. produced the recombinant proteins and conducted SPR. M.R.W., T.G., and J.K. conducted the Cryo-Em analysis. A.M.K., A.M., Y.M., C.Y. and L.M.-S. conducted the *in vitro* and *in vivo* live virus assessments. P.A.G. and N.B.E. acquired the clinical specimens. D.D.H., L.L., and Q.W. conducted the pseudovirus neutralization assay. J.J.K., L.M.-S., M.R.W., and D.D.H. supervised the work. M.S.P., J.J.K., L.M.-S., and M.R.W. wrote the manuscript. All authors reviewed the results and approved the final version of the manuscript.

