## Supplemental Figure 1 for "Potent neutralization by a receptor binding domain monoclonal antibody with broad specificity for SARS-CoV-2 JN.1 and other variants"

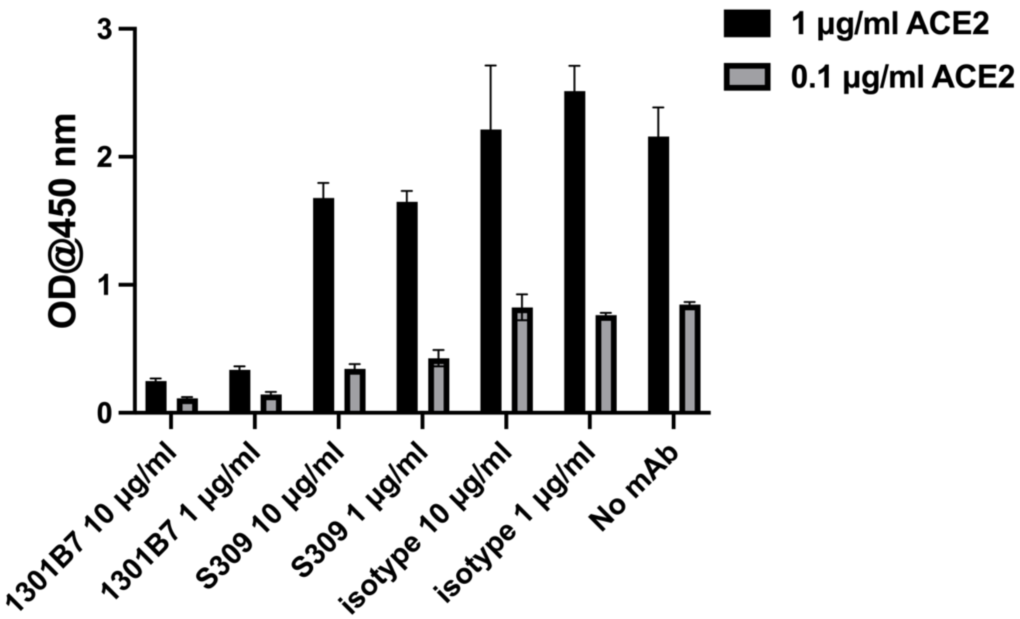


**Supplemental Figure 1. 1301B7 competes with human ACE2 for SARS-CoV-2 RBD binding.** Binding of 0.1 or 1.0 µg/ml of streptavidin-conjugated recombinant human ACE2 to SARS-CoV-2 RBD in the presence or absence (no mAb) of indicated mAb was determined by ELISA.
