## Supplemental Figure 2 for "Potent neutralization by a receptor binding domain monoclonal antibody with broad specificity for SARS-CoV-2 JN.1 and other variants"

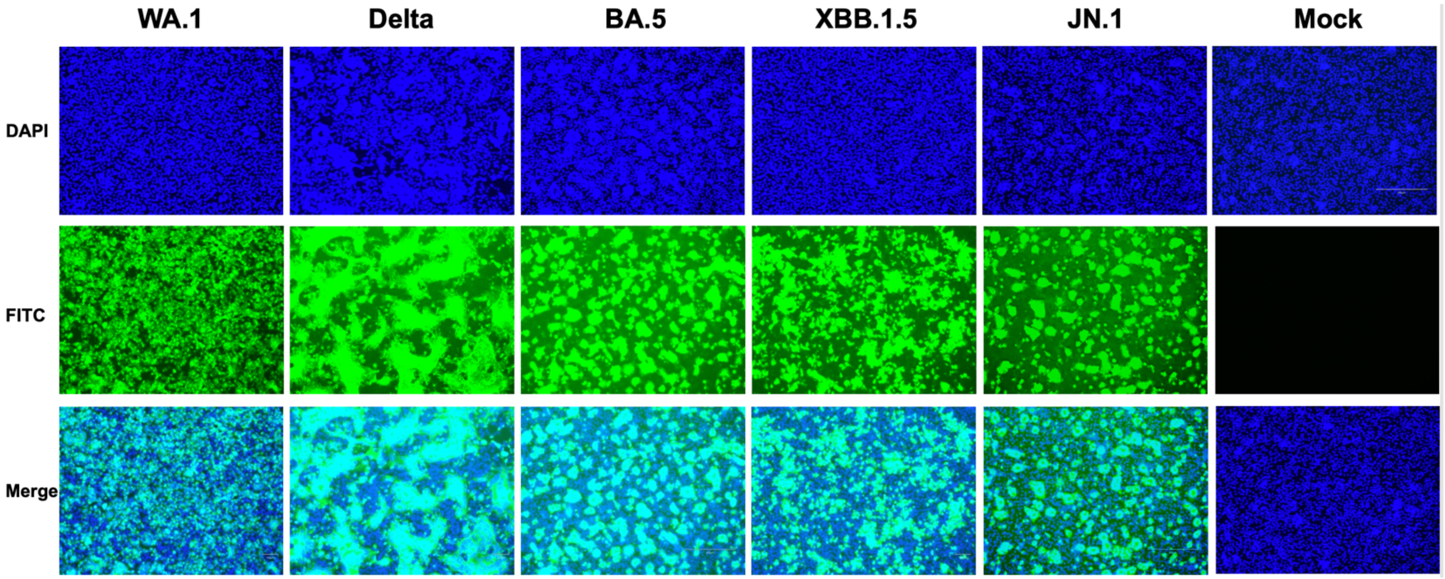


**Supplemental Figure 2. 1301B7 binds to SARS-CoV-2 infected cells.** Confluent monolayers of Vero AT cells were infected (MOI 0.1) with SARS-CoV-2 WA-1, Delta, Omicron BA.5. Omicron XBB1.5, or Omicron JN.1. Mock-infected cells (right) were included as control. Cells were incubated with 1301B7 hmAb (1 µg/ml) and developed with a FITC-conjugated secondary anti-human Ab. 4,6-Diamidino-2-phenylindole (DAPI; blue) was used for nuclear stain. Scale bar indicate 100 µm.
