## Supplementary figures and images for "Potent neutralization by a receptor binding domain monoclonal antibody with broad specificity for SARS-CoV-2 JN.1 and other variants"

### Supplemental Figure 3

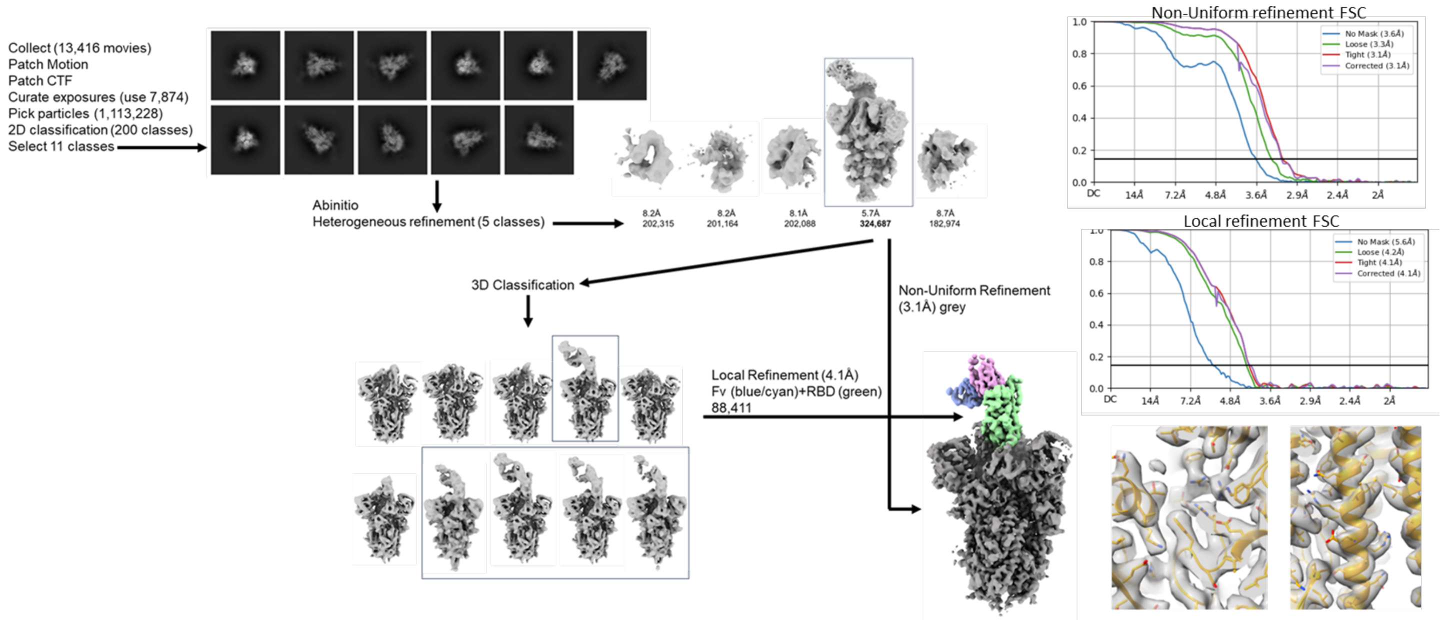


**Supplemental Figure 3. Flow diagram for EM data processing.**
